# Spatial redistribution of neurosecretory vesicles upon stimulation accelerates their directed transport to the plasma membrane

**DOI:** 10.1101/2021.05.20.444910

**Authors:** Elaine B. Schenk, Frederic A. Meunier, Dietmar B. Oelz

## Abstract

Through the integration of results from an imaging analysis of intracellular trafficking of labelled neurosecretory vesicles in chromaffin cells, we develop a Markov state model to describe their transport and binding kinetics. Our simulation results indicate that a spatial redistribution of neurosecretory vesicles occurs upon secretagogue stimulation leading vesicles to the plasma membrane where they undergo fusion thereby releasing adrenaline and noradrenaline. Furthermore, we find that this redistribution alone can explain the observed up-regulation of vesicle transport upon stimulation and its directional bias towards the plasma membrane. Parameter fitting indicates that in the deeper compartment within the cell, vesicle transport is asymmetric and characterised by a bias towards the plasma membrane. We also find that crowding of neurosecretory vesicles undergoing directed transport explains the observed accelerated recruitment of freely diffusing vesicles into directed transport upon stimulation.

## Introduction

Neuroendocrine chromaffin cells in the adrenal medulla are a model system to study the exocytosis of secretory vesicles (SVs) which is preceded by their transport from the Golgi apparatus to their site of exocytosis on the plasma membrane [1]. Defects in this process have been tied to a range of neurodegenerative diseases [2] by mechanism that are not yet fully elucidated [3]. The highly crowded and stochastic nature of the cytoplasm, however, represents a major difficulty in exploring intracellular transport systems and as a result many of the mechanisms involved in bringing vesicles towards the plasma membrane remain elusive [1].

Microtubules and actin filaments perform a vital role in the transport of secretory vesicles and can be considered the tracks of the intracellular transport network [4]. They are semi-flexible and some of them can be several μm in length.

In the central region of the cytoplasm both systems of cytoskeleton filaments, F-actin and microtubules, are involved in the cytoplasmic transport of secretory granules [5–7] through molecular motor proteins. In this cellular region microtubules are mostly aligned radially while close to the cortex their density is lower and they align tangentially with the cortex [6]. In the periphery of the cell, transport along the actin filament network occurs through processive molecular motors such as myosin-V [8, 9] [1, 10] and non-processive such as Myosin-II [11]. Recruitment of SVs to the cortical actin network occurs via the action of Myosin-VI short-insert isoform which has both a processive and a non-processive function [12].

The sketch shown in Fig 1 shows the cytoplasm of chromaffin cells with neurosecretory vesicles immersed in and occasionally transported along microtubules (thick lines) or actin filaments (thin lines). In addition to its role as a barrier between the cytoplasm and the active sites [13, 14] the cortical actin network has an active role in exocytosis. It contributes to the transport of SVs in the periphery of the cell [15], facilitates the transport of SVs through the cellular cortex towards the sites of exocytosis and it also contributes mechanically to the exocytosis [16, 17].

**Fig 1.**
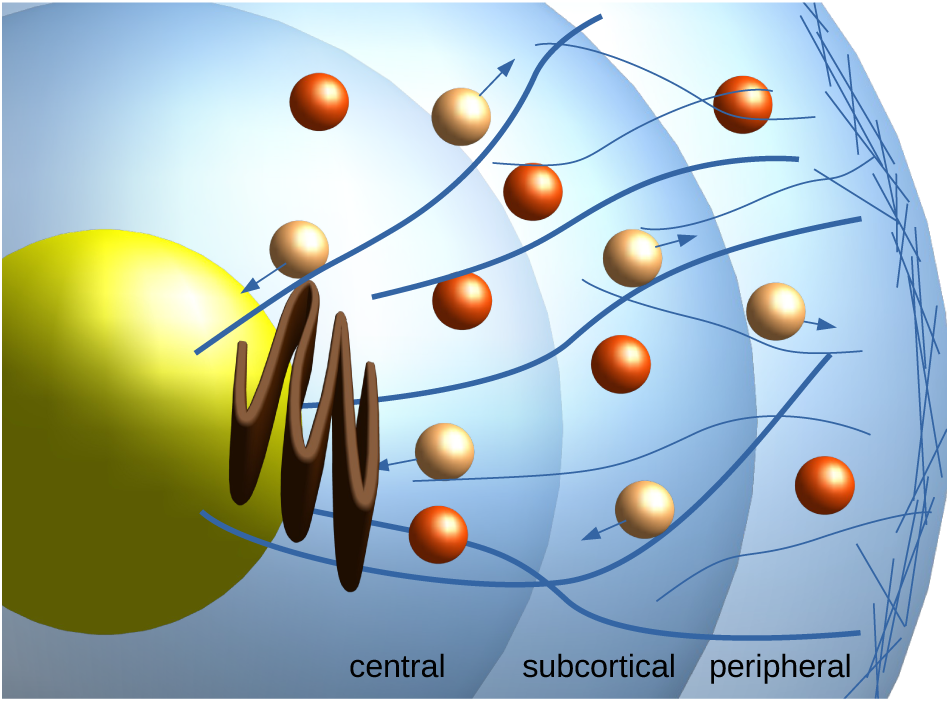
Sketch of secretory vesicle transport between the Golgi apparatus (brown) and the cortical actomyosin network in chromaffin cells. Microtubules are shown as thick blue curves, actin filaments as thinner fibres. Secretory vesicle currently undergoing transport are represented by the lighter coloured spheres with arrows denoting the direction of transport whereas the darker orange spheres represent vesicles that are freely diffusing in the cytoplasm.

Vesicles undergoing directed motion along microtubules often cover relatively large distances [18]. Some vesicles exhibit a restricted behaviour [19] termed “caged”, which is generally associated to anchoring to the actin meshwork [12]. It has also been suggested that within the dense actin meshwork, secretory vesicles can become effectively trapped and hence also display caged behaviour [18].

In [19] Live-cell imaging of nicotine stimulated bovine adrenal chromaffin cells was used to track the motion of secretory vesicles in 3-dimensions by confocal microscopy. Vesicles were categorised as undergoing either directed, free or caged motion and according to their distance from the plasma membrane. The measured transition rates implied that upon nicotine stimulation, cells spatially adjust their secretory vesicle pools and that upon stimulation both the fraction of vesicles undergoing directed transport increases as well as the rate by which vesicles enter directed transport – both contributing to the efficient replenishment of the pools of releasable vesicles. What feedback mechanisms, however, trigger those adjustments is not currently known.

The aim of this study is to introduce a quantitative description of secretory vesicle transport in chromaffin cells that is well adapted to the available data. Our goal is to show that spatial redistribution of vesicles can trigger the observed up-regulation of directed transported upon stimulation.

Modelling studies of intracellular transport have so far focused on transport along and within bundles of fibres [20–23]. On the other hand cytoplasmic vesicle transport as part of the secretory pathway has not received much attention from modellers so far: In [24] the biochemical reaction network underlying exocytosis has been modelled through a system of rate equations. Agent based models have been considered in various studies such as in [25] which focuses on simulation methods, as well as in [26] on spatial aspects of vesicular sorting into different compartments and in [27] on aspects of pattern formation.

Mathematical modelling of the transport of secretory vesicles in chromaffin cells has so far been focusing on arrival and release statistics. Amperometric techniques make it possible to record the discrete arrival times of newly trafficked vesicles at the plasma membrane and hence provide insight into the underlying nature of the secretory pathway. In [28] it was shown that these arrival times follow a non-Poissonian probability distribution suggesting that there is an active recruitment process underlying the replenishment of releasable vesicles upon stimulation. More specifically, through the coupling of an attractive harmonic potential to a random process modelling vesicle migration, key aspects of arrival time statistics could be explained. Nevertheless, such a phenomenological model does not have the potential to link this effect to structural properties of the intracellular environment.

In order to address this issue, a recent study [29] suggested a system of advection-diffusion equations [30] as a tool to explore the various features of up-regulation of directed vesicle transport reported in [19]. It was found that spatial redistribution triggered by nicotine stimulation explains the observed bias toward *outward transport as compared to inward* transport upon stimulation. The parameter space for this class of models, however, turned out to be too high-dimensional prohibiting meaningful parameter fitting given the limited amount of available data [19]. As a consequence, this study could not address the question whether and under which additional assumptions the spatial redistribution that occurs upon stimulation can explain other more intricate aspects of up-regulation such as the much larger overall fraction of vesicles undergoing directed transport and the accelerated recruitment into directed transport.

In our study we construct a conceptually simpler Markov state model to address these questions. This allows for a direct incorporation of the transition rates which are reported in [19] individually for each of the three spatial compartments: central, subcortical and peripheral.

We indeed find that the Markov chain based on the reported transition rates predicts spatial redistribution upon stimulation. Adding the additional assumption obtained through parameter fitting that the transport network in the central part of the cell has an outward bias, we find that this amplifies the overall fraction of vesicles undergoing directed transport in agreement with observations. We illustrate the underlying mechanism using an even simpler minimal Markov state model.

We also find that spatial redistribution accelerates the transition rates into directed transport as reported in [19] provided that the capacity of transport fibres is limited.

### Preliminary experimental results

In [19] bovine adrenal chromaffin cells were imaged using time-lapse z-stack confocal imaging. A number of equatorial slices in the x-y plane of chromaffin cells were imaged so that 20% of the cell’s latitudinal range was covered. As a consequence, most of the secretory vesicle activity occurred within the imaged region. Individual vesicles were then tracked and the growth of each trajectory’s mean square displacement as a function of time - sub-linear, linear or supra-linear - was used to categorize them as either *caged*, *free* or *directed*. Comparing vesicles in control and in nicotine stimulated cells, the following key findings were reported:

1. *Up-regulation of directed transport*: Upon stimulation the fraction of vesicles undergoing directed motion grows from roughly 38% to 52% (Fig 1E in [19] which is reproduced in Fig 2E).
2. *Bias towards outward transport:* While vesicle transport is unbiased in control cells, in stimulated cells the ratio of outward vs. inward transport is roughly 2:1 (Fig 3S and 3T in [19]).

**Fig 2.**
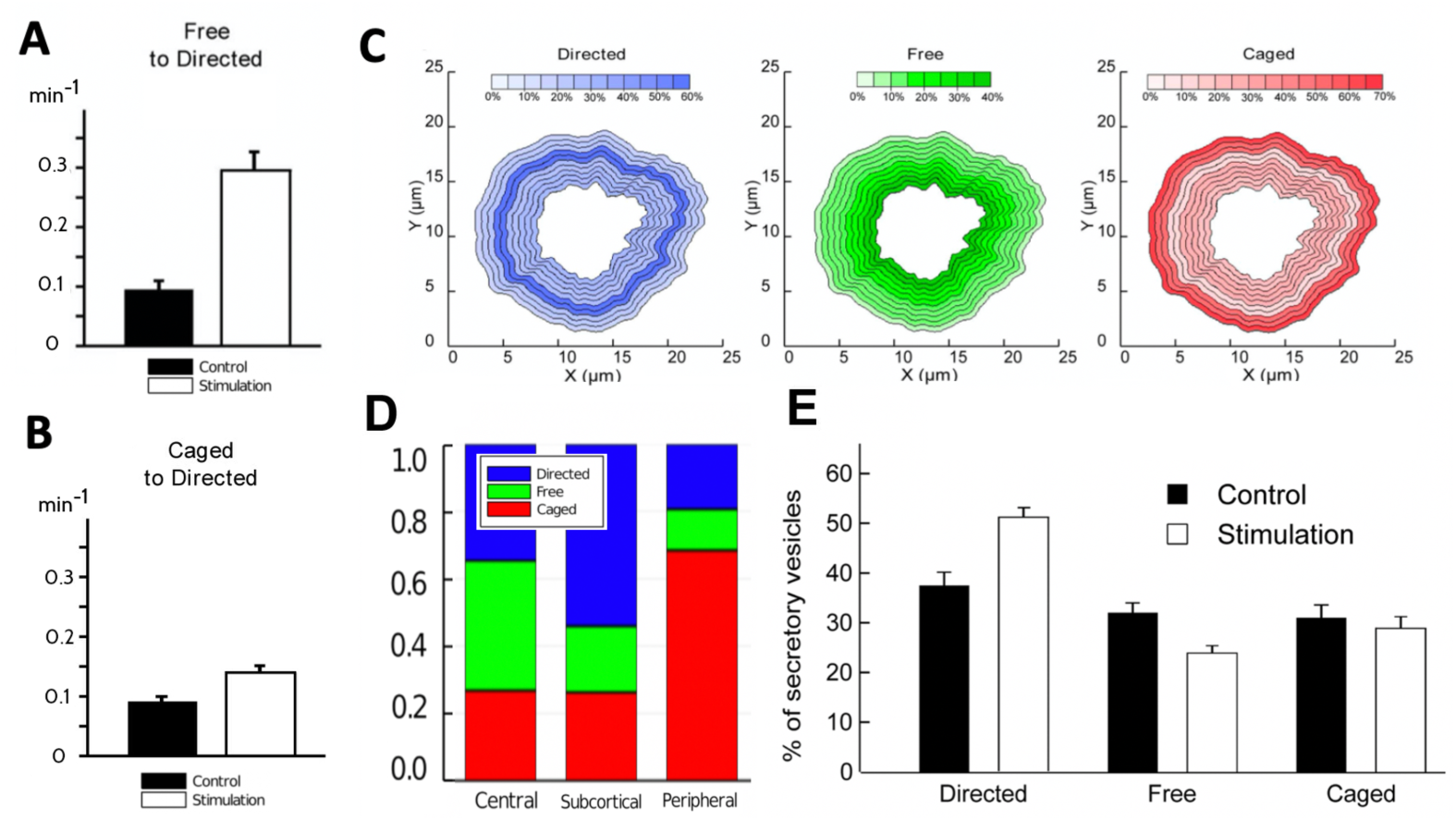
Statistics of tracked vesicles reported in [19]. (**A**) Figures 3H (with rates per 2 min) in [19]: average transition rates from free to directed motile state in control and stimulation. (**B**) Figures 3B (with rates per 2 min) in [19]: average transition rates from caged to directed motile state in control and stimulation. (**C**) Figure 2C in [19]: Proportions of vesicle motile behaviour in spatial bands of radial width 0.5 μm. (**D**) Data in Fig 2C from [19] averaged within the three spatial compartments. (**E**) Figure 1E in [19]: Percentages of secretory vesicles in each motile state in control and stimulation.

**Fig 3.**
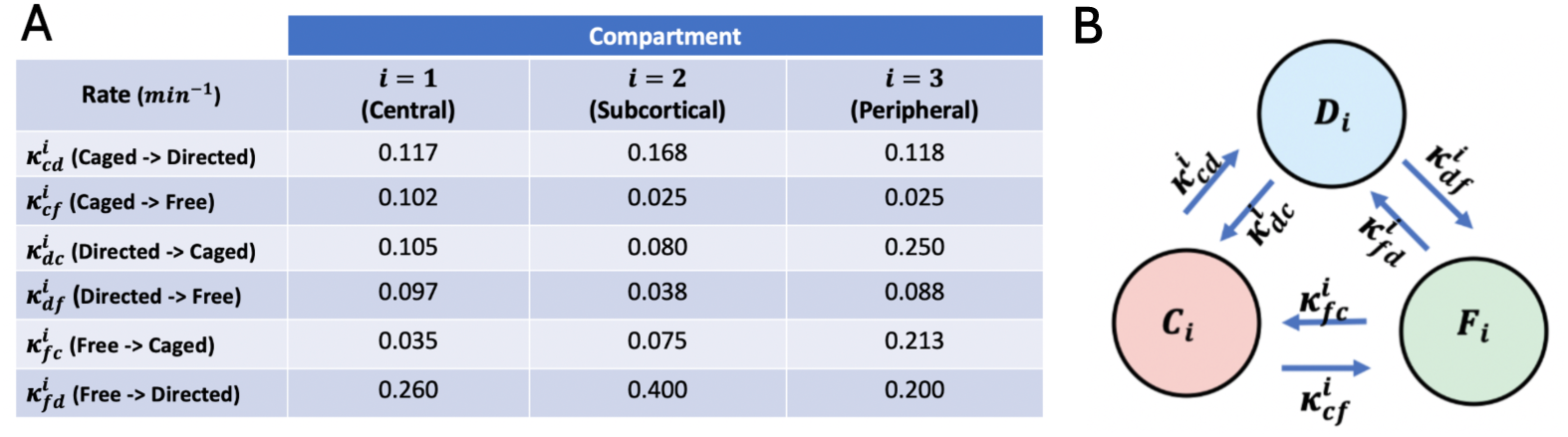
(A) Experimentally observed transition rates. Table of transition rates reported in [19]. (**B**) Graphical sketch of 3-state Markov chain model (*C_i_*(aged), *D_i_*(irected), *F_i_*(ree)) for single spatial compartments *i* = 1, 2, 3.

In addition to categorising vesicle trajectories, the transitions of vesicles from one motile state to another were recorded in trajectories spanning periods of 2 minutes. One of the key observations was that

3. stimulation leads to a *three-fold increase of the rate by which vesicles transition from free (unbiased random walk) motion to directed motion* (Fig 3H in [19] reproduced in Fig 2A).

Moreover, determining the transition rates within spatial bands of radial width 0.5 μmrevealed three functionally distinct ring-shaped spatial compartments (visualised in Figure 1): the *peripheral* (0.5-1.5 μm distance from cell membrane), *subcortical* (1.5-2.5 μm) and *central* (2.5 - 5.0 μm). Note that when determining the average transition rates for these compartments we omit the outermost radial band right at the cortex which is mostly populated by caged vesicles. Fig 3 provides the transition rates reported in Fig.3J-R in [19] (per 2 minutes) converted to numerical values per 1 minute as well as a sketch of the underlying 3-motile-states model.

They provide at least speculative insight into the underlying architecture of the intracellular transport network: The higher free-to-directed transition rate in the subcortical compartment suggests the presence of a higher density of cytoskeleton fibres in this region. Another observation is that the free-to-caged transition rate is significantly elevated in the peripheral region which might reflect a relatively higher density of the peripheral actin meshwork [31]. The higher caged-to-directed transition rate in the subcortical compartment as compared to the practically identical rates in the central and peripheral ones suggests that the microtubules and actin meshwork are both dense in this compartment or that they achieve significant overlap.

The directed-to-free transition rates are relatively higher in the central and peripheral compartments in comparison to the subcortical rate. We hypothesise that this is due to the higher number of microtubules terminating within these regions by which the faster off-rates there might also reflect the detachment of vesicles after reaching the “end of the track”. Another interesting observation from Fig 3 is that the caged-to-free transition rate is relatively higher in the central compartment, whereas free-to-caged rate is the lowest in the central compartment. This might be evidence that recently synthesised vesicles might enter the transport network as initially caged vesicles which is supported by the literature [32].

## Materials and methods

Our goal is to develop a Markov state model based on the transition rates listed in Fig 3. In [19] these have been published for stimulated cells, but not for control cells. We will therefore assume that the transition rates listed in Fig 3 in principle are valid irrespective of stimulation vs non-stimulation. Note that in the context of our model, this implies that stimulation does not structurally alter the characteristics of how secretory vesicles behave in the cytoplasm, and especially interact with the cytoskeleton.

We will show that such a model is able to explain the key experimental observations listed above. Spatial inhomogeneity of the cytoskeleton, specifically the non-uniform spatial distribution of outward vs inwards directed vesicle transport, coupled with the spatial redistribution of vesicles upon stimulation, will trigger up-regulation of outward transport and of overall transport as listed above. We refer to this mechanism as the “acceleration through spatial redistribution” hypothesis. We will start by introducing a minimal (“toy”) model in order to provide an intuitive initial illustration of the mechanism.

### Minimal 4-state model

The following 4-state continuous time Markov chain visualised in Fig 4 provides a proof of concept for the “acceleration through spatial redistribution” hypothesis.

**Fig 4.**
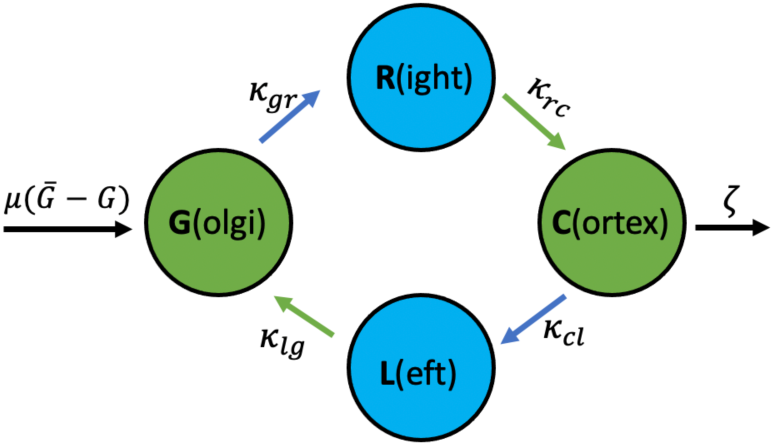
Graphical sketch of the minimal 4-state model.

In this simple model vesicles exist in one of four states, respectively spatial compartments: They are generated at the Golgi apparatus (G), from where they might engage in outward – visualised as right-directed – transport (R) towards the cell cortex (C). Vesicles at the cortex engage in in-ward – shown as left-directed (L) – transport and ultimately arrive back at the Golgi. Alternatively, vesicle at the cortex may enter the pathway towards exocytosis with rate *ζ*. This pathway acts as a sink whereas the cell’s synthesis of vesicles near the Golgi represents a source of new secretory vesicles. We assume that vesicle synthesis is regulated by feedback mechanisms which aim at maintaining a given number of 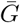 vesicles at the Golgi apparatus.

This gives rise to the system of rate equations

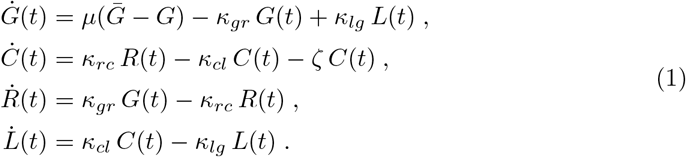

In order to verify the up-regulation through spatial redistribution hypothesis, we take *ζ* as zero in a control cell and assume that stimulated cells are characterised by *ζ* > 0. In our assessment we only consider the steady state solution (for details see supplementary section S1 Steady state solution of the minimal 4-state model) where *μ* is large, i.e. where 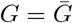 and

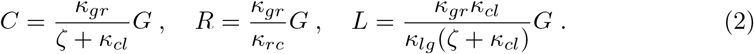

In the context of this minimal model up-regulation of outward transport is characterised by how 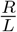 depends on *ζ*, namely

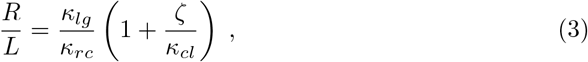

which increases monotonically as a function of *ζ* since all transition rates are positive. This shows that the minimal model predicts a bias towards outward transport in response to stimulation. Note that the decrease of the number of vesicles in *L* is triggered by a preceding decrease in the compartment C (see (2)). In this sense the observed up-regulation is a consequence of the adjustment of the spatial distribution upon stimulation.

Note that to mimic a ratio of *R* : *L* ∼ 1 which is observed in control cells (*ζ* small) the rates *κ_lg_* and *κ_rc_*, which both represent off-rates from vesicle transport, are required to be approximately equal as a consequence of (3).

Up-regulation of directed transport upon stimulation, on the other hand, is characterised by the ratio 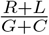 which – at steady state – is given by

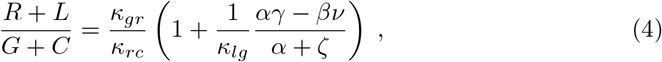

where 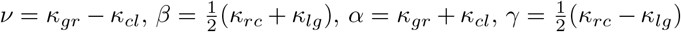. Note that *α* and *β* are both positive. Therefore, the minimal model predicts up-regulation of directed transport only if *ν >* 0 and/or *γ <* 0. The fact that we need *κ_rc_* ∼ *κ_lg_* for an 1:1 ratio of right and left moving vesicles in control suggests *γ* ∼ 0 and therefore we require *ν >* 0.

Interpreting the on-rates *κ_gr_* and *κ_cl_* as representatives of effective fibre densities - the more cytoskeleton fibres and motor proteins moving in the right direction are available, the larger the respective on-rates – the result that *ν >* 0 indicates that there should be a higher net density of fibres triggering outward transport as compared to fibres resulting in transport away from the cortex. It is worth noticing that small *κ_cl_* favours both, up-regulation of directed transport in (4) and a strong bias towards outward transport (*R/L*, see (3)) upon stimulation.

Finally, it is important to note that there is an inherent asymmetry in this minimal model. Specifically, only transitions from the spatially interior state G to right directed motion are considered and not vice versa. Similarly, we only consider transitions from the spatially outward state C to left directed motion and not vice versa. This restriction makes sense for a simplified model thought to be coarse-graining a more intricate underlying system. In the following development of the full Markov state model we consider three spatial compartments with bi-directional transitions modelling vesicle attachment and detachment from cytoskeleton fibres and molecular motors. The asymmetry of the minimal model will be replaced by transitions between spatial compartments modelling directed transport.

### Markov state model

Our goal is to formulate a mathematical model which explains the key experimental observations reported in [19]. To this end we refine the minimal model visualised in figure 4 by adding spatial compartments building on the measured rates listed in Fig 3.

We represent the spatial distribution of vesicles by assigning them to one of the three compartments *central* (i=1), *subcortical* (i=2) and *peripheral* (i=3), which mimics the terminology used in [19].

In each spatial compartment vesicles are either in state *F_i_* (diffusing *freely* in the cytoplasm), *C_i_* (*caged*, i.e. undergoing random, though spatially restricted motion) or undergoing directed motion in state *R_i_* (towards the cell periphery usually visualised on the *right*) or *L_i_* (towards the cell centre usually visualised on the *left*).

Figure 5A provides a visualisation of the transitions occurring within the individual spatial compartments of the model. Note that the experimental study only provides information about the rates by which either free or caged vesicles engage in directed transport, but not about the direction of transport (Fig 3).

**Fig 5.**
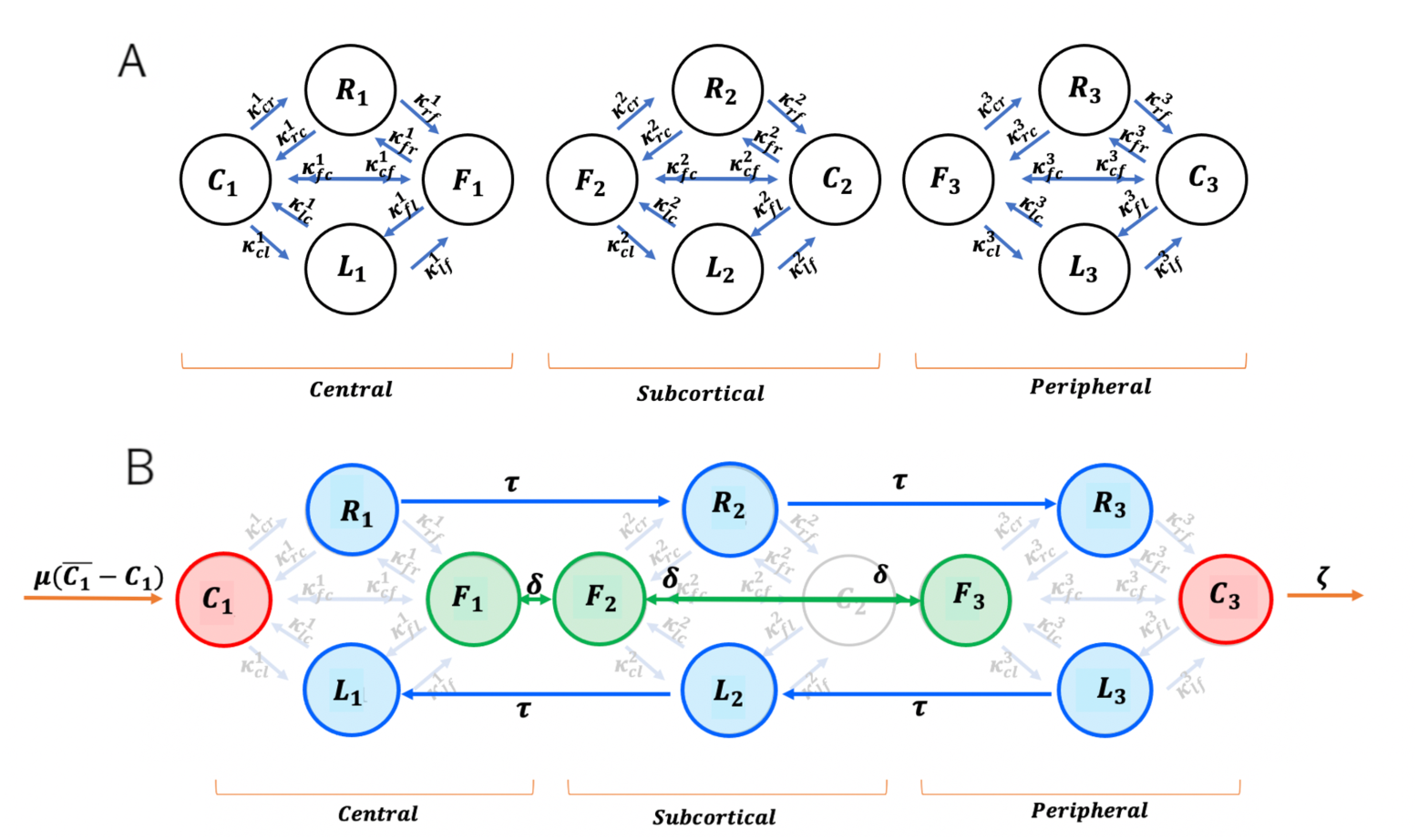
Visualisation of Markov states with intra-compartmental transitions (A) and inter-compartmental transitions (B).

For this reason we introduce the structural parameter 0 ≤ *ρ_i_* ≤ 1 for each of the three spatial compartments *i* = 1, 2, 3. The role of this parameter is to determine how the experimentally recorded on-rates into directed transport 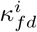 and 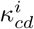 from figure 3 distribute into outward (right) and inward (left) transport. Using the structural parameters we write the transition rates from the free and caged states into either left or right transport shown in Fig 5A as

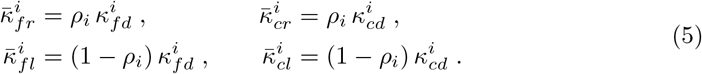

Here we use the overbar notation to indicate that these rates will be further modified below. One way of interpreting the factors *ρ_i_* is as the fraction of fibres within one of the spatial compartments along which vesicles will be transported outwards vs inwards. This implies that *ρ_i_* does not reflect the concentration of fibres within the respective spatial compartment. Instead the rates 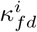 are likely to reflect the net fibre densities in any given compartment.

Similar considerations for off-rates are not necessary, so we assume that

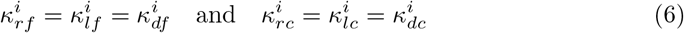

for all three spatial compartments *i* = 1, 2, 3, i.e. the experimentally observed off-rates from directed transport reported in figure 3 taken from [19] are used irrespective of the direction of transport.

In order to connect the so-far isolated spatial compartments sketched in Fig 5A, we introduce additional transitions accounting for the random and directed motion of vesicles. The mean square displacement per time of free vesicle trajectories in 3D has been determined in [19] (Fig S1) as 0.75 μm^2^/min. Assuming an average distance of 1.7 μm between our three, linearly aligned, spatial compartments this suggests a inter-compartment transition rate of *δ* = (0.75/6)/1.7^2^ min^−1^ ≈ 0.04 min^−1^ to account for random motion of free vesicles. The corresponding transitions between the states of free vesicles in neighbouring compartments are visualised in Fig 5B as bidirectional arrows.

We also include uni-directional transitions between the directed states in order to represent directed transport of secretory vesicles. In Fig 5B these are shown as blue arrows. Instantaneous maximal vesicle speeds of up to 0.05 μm s^−1^ have been reported in [19]. This would suggest transition rates of 0.05 × 60/1.7 ≈ 1.8 min^−1^. In steady state solutions of the Markov state model, however, these fast transition rates would result in vesicles pooling at the states *R*_3_ and *L*_1_ which is not supported by observations [19]. Instead, we assume that vesicles achieve maximal speeds only intermittently and estimate the transition rate of vesicles undergoing directed transport as *τ* = 0.15 min^−1^.

Additional transitions embed the Markov State model (visualised in Fig 5B within the life-cycle of secretory vesicles. We model vesicle synthesis as an in-flux of vesicles into *C*_1_ (the caged pool close to the Golgi apparatus) with rate 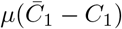. At steady state this term effectively imposes the constant value 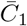 on the number of vesicles in *C*_1_, so for our purposes *μ* is an arbitrary large value which has no effect on the steady state solutions and 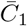 is an arbitrary constant which factors out of the steady state solution after we normalise either by the total number of vesicles in the system or in the respective spatial compartment. Having vesicles enter the system through the caged node *C*_1_ as opposed to the central node of free vesicles *F*_1_ reflects our hypothesis (supported by [32]) that vesicles travel through an actin meshwork adjacent to the Golgi upon entering the system.

Finally, as in the minimal 4 state model, we model the onset of the pathway through which vesicles undergo exocytosis by an outgoing-transition (rate) from the pool of caged vesicles in compartment *i* = 3 (peripheral) denoted by *ζ*. This reflects the role the actin cortex plays in the exocytosis pathway [31]. With these parameter values, especially with the estimated values for *τ* and *ζ* steady state simulations of stimulated cells exhibit relative shares of vesicles (free, caged and undergoing directed motion) Fig 7A which coincide well with measured values from [19] visualised in Fig 2C and D. Note that the full system of rate equations corresponding to the Markov chain shown in Fig 5 is shown in supplementary section S2 Governing Equations.

**Fig 6.**
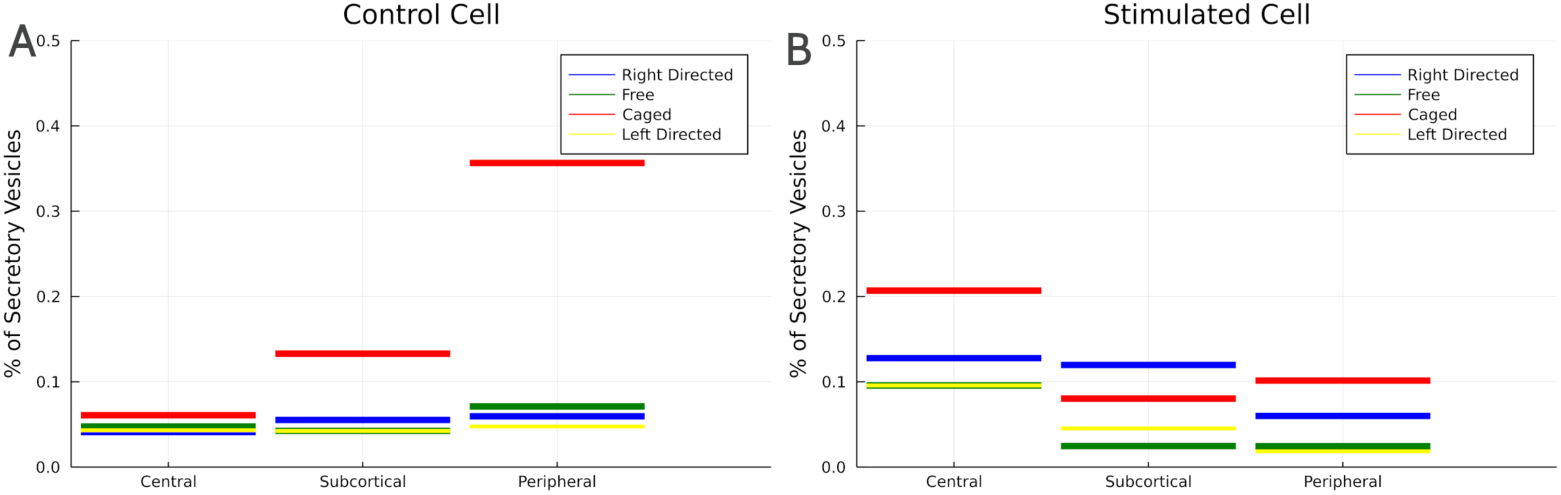
Simulated steady state probability distributions of vesicles in control (A) and upon stimulation (B), both normalised so they admit a probabilistic interpretation.

**Fig 7.**
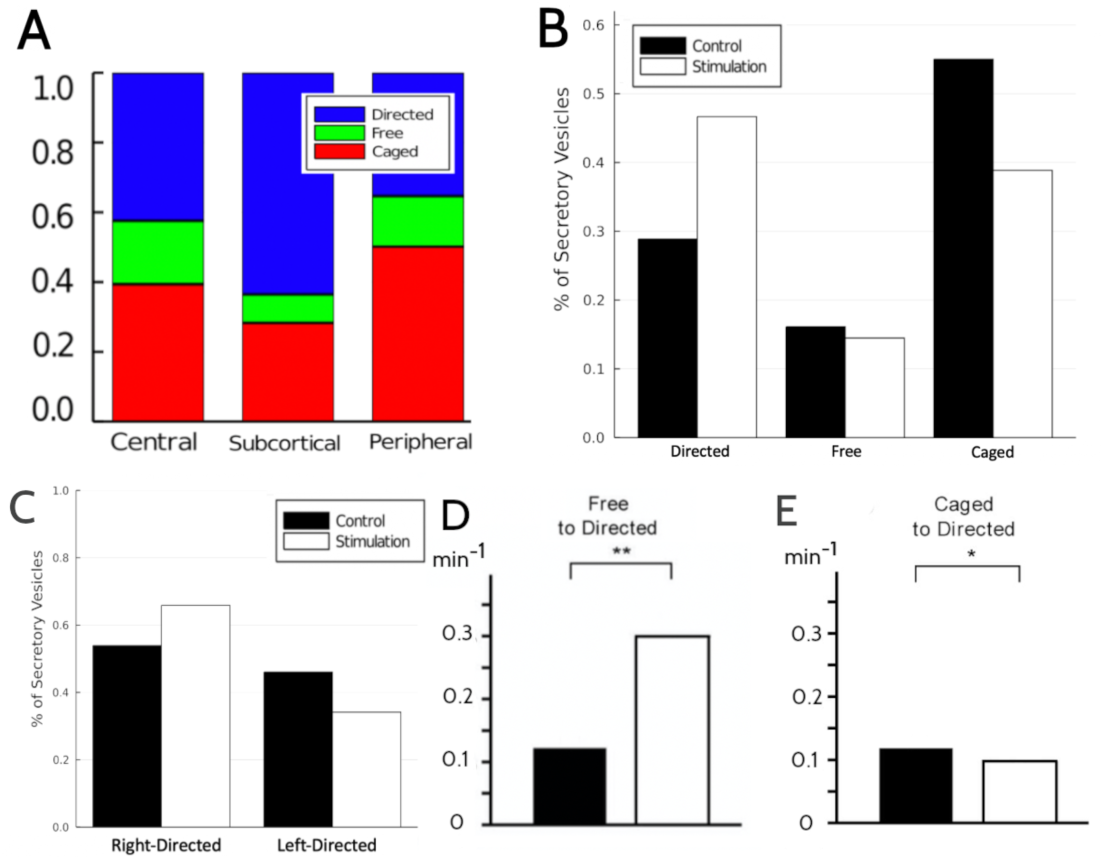
Steady state simulation results: **A** Percentages of secretory vesicles in each motile state (stimulation). **B** Percentages of secretory vesicles in each directed (right vs left).**C** Intra-compartmental proportions in stimulated cells. **D** Aggregated transition rates from free to directed motion. **E** Aggregated transition rates from caged to directed motion.

### Fitting of structural parameters

We determine the structural parameters *ρ*_1_, *ρ*_2_ and *ρ*_3_ (see Table 1) by fitting (least-squares) the predicted ratios of right vs left moving vesicles denoted by Φ*_ζ_* and the predicted ratio of directed motion vs random motion denoted by Ψ*_ζ_*. These quantities are given by

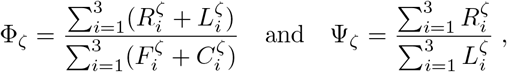

where i is the spatial compartment index and 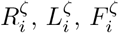 and 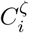 are the steady state abundances for a given exocytosis rate *ζ*.

**Table 1.** Parameter values

| Description | Symbol | Value | Reference |
| --- | --- | --- | --- |
| Inter-compartmental transition of free vesicles | $\delta$ | $0.04 \text{ min}^{-1}$ | Figure S1 in [19]. |
| Transition into exocytosis pathway (control) | $\zeta_{\text{ctr}}$ | $0.0 \text{ min}^{-1}$ | Estimated |
| Transition into exocytosis pathway (stimulated cells) | $\zeta_{\text{stim}}$ | $0.15 \text{ min}^{-1}$ | Estimated |
| Directed spatial transition | $\tau$ | $0.15 \text{ min}^{-1}$ | Estimated |
| Outward vs inward transport | $\rho_1, \rho_2, \rho_3$ | 0.75, 0.6, 0.4 | Estimated (see supplementary section Fitting of structural parameters) |
| Repression (crowding) coefficients | $\hat{R}_i, \hat{L}_i$ ($i = 1, 2, 3$) | 0.2 | Estimated |

We take into account the deviations of those quantities from the observed ones for both, stimulated cells and control cells. The error functional which we minimise is therefore given by

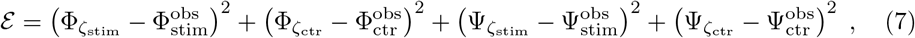

where the experimentally observed values are given by 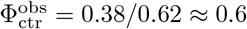, 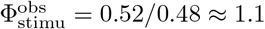 (see Fig 2E) as well as 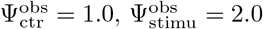.

### Modelling of carrying capacities

Steady state simulation results with the constant transition rates (5), however, are not consistent with the threefold increase in the average free-to-directed transition rate observed upon stimulation (Fig 2A). Spatial redistribution alone cannot account for this since the slowest - among all spatial compartments – reported transition rate 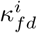 in stimulation is 0.2 min^−1^ (Fig 3), whereas the average rate reported in control is 0.1 min^−1^ (Fig 2A, which reports rates *per 2 min*).

We hypothesise that the observed increase in the average free-to-directed transition rate upon stimulation is a consequence of the global decrease in vesicle abundance occurring at stimulation. We therefore suggest that in control transitions are slowed down by crowding of vesicles undergoing directed transport.

We use Hill functions (for a repressor [33]) to model the limiting effect of carrying capacities on all other transition rates into directed transport by the following correction factors

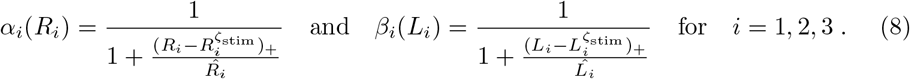

The Hill functions are parametrised by the repression coefficients 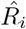 and 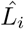 (*i* = 1, 2, 3). To ensure that this modification does not alter the steady state population sizes in stimulation, 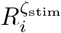 and 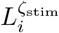, we include these values in the correction factors (8) taking only the positive part of 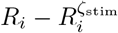 and 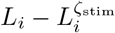, respectively.

It obviously appears problematic to include steady state abundances in stimulation in the correction factors. To remove these dependencies one might rewrite the correction factors as a classical Hill function with two constants, namely *α_i_*(*R_i_*) = *μ_i_/*(1 + *R_i_/ν_i_*)(for 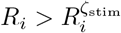) and analogous expressions for *β_i_*. Nevertheless, such rewriting of the correction is not required to run the simulations and it would render our choice of parameters less transparent. We therefore keep the notation used in (8).

Note that applying the correction factors (8) to the transition rates (5) would also slow down transitions from the pool of caged vesicles into transport states. Reported transition rates from caged into directed motion (averaged among spatial compartments), however, only exhibit minor variation between control and stimulation (Fig 2A). We argue that for caged vesicles the speed-up in response to less crowding along transport fibres upon stimulation might be compensated by faster turnover of the actomyosin cortex upon stimulation [31] which could prevent a large fraction of caged vesicles from binding to fibre-motor-protein complexes.

As a consequence we keep the peripheral caged-to-directed transition rates unmodified, while rescaling all rates listed in (5) as follows

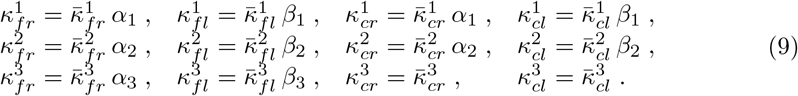

In the absence of more detailed data, and since we only seek to provide a preliminary proof of concept for the idea that crowding can explain the 3-fold speed-up of transitions from free to directed (Fig 2A) we choose a single constant for the repression coefficients (see Table 1).

Also note that to keep the fitting procedure simple we do not reassess the structural parameters *ρ_i_*, *i* = 1, 2, 3 with the modified transition rates. Indeed they only have a minor effect on the capacity of the model to correctly predict up-regulation of outward and total transport as shown below in section Spatial redistribution can explain both, up-regulation of directed transport and bias towards the plasma membrane.

## Results

### Stimulated exocytosis triggers spatial rearrangement of chromaffin vesicles

We numerically compute the steady state solutions of the Markov state model visualised in Fig 5 with the rates (9) and (6).

In simulated control cells, the steady state distribution of vesicles (Fig 6A) is characterised by a large portion of vesicles (of about 38%) being pooled in the caged state at the cortex. In the corresponding steady state distribution for a stimulated cell (Fig 6), this pool of vesicles ready to enter the pathway leading to exocytosis is largely reduced. This results in caged vesicles being distributed more evenly between the three spatial compartments upon stimulation with most of them now being located in the central region of the cell. All together this accounts for a significant shift in the spatial distribution of vesicles from the cell periphery to the cell centre as a consequence of the permanent out-flux of vesicles at the cortex in response to secretagogue stimulation.

From the results of Fig 6 we extract the intra-compartmental proportions for a stimulated cell which are visualised in Fig 7A and qualitatively agree with the experimentally recorded intra-compartmental proportions shown Fig 2C and D.

### Spatial redistribution can explain both, up-regulation of directed transport and bias towards the plasma membrane

Our model predicts an increase from 29% to 47% in the share of vesicles undergoing directed transport in stimulated cells vs control (Fig 7B). This compares well with the experimental data reproduced in Fig 2E. Likewise the rate of outward moving vesicles vs. inward moving vesicles increases from about 1:1 in control to about 2:1 in stimulated cells (Fig 7C) which also coincides with the data reported in [19].

The vesicle distributions in Fig 6 reveal that upon stimulation the relative abundance of vesicles moving outwards (blue) increases in the central and subcortical compartment. On the other hand the relative abundance of vesicles moving inwards (yellow) decreases in the peripheral compartment, though to a lesser extent. Both effects contribute to the up-regulation of total and outward directed transport. They are both triggered by spatial redistribution of vesicles upon stimulation since the states in which most of the redistribution takes place are those which precede states of direct motion: from *C*_1_ and *F*_1_ vesicles transition into *R*_1_ – and later *R*_2_ and *R*_3_, from *C*_3_ they transition into *L*_3_ – and later back into *L*_2_ and *L*_1_.

This mechanism is overshadowed by the dense network of potential transitions in the Markov State model Fig 5 and so it is easier to understand it in the context of the minimal model Fig 4. This simpler model also features redistribution of non-moving vesicles upon stimulation, namely through the number of vesicles in state C(ortex) which decreases upon stimulation (*ζ* large) according to Eq (2). As a consequence, upon stimulation the number of vesicles in the state L(eft) also decreases, since C is the only state from which vesicles transition into L. This readily explains the increase of the ratio R:L under stimulation (see Eq (3)) in the minimal model. It also indicates that in the full Markov state model the mechanism which drives the increase of the ratio of outward vs inward transport upon stimulation is the redistribution of caged and free vesicles upon stimulation.

The mechanism which drives the up-regulation of the total share of vesicles undergoing directed transport is more obscure. For the minimal model we have concluded that up-regulation of total directed transport requires *ν = κ_gr_* – *κ_cl_ >* 0 in Eq (4) (or alternatively *γ <* 0 which we ruled out assuming that 2*γ = κ_rc_ – κ_lg_* = 0). In other words, the rate by which vesicles enter the state representing transport to the R(ight) should be faster than the corresponding rate by which vesicles enter the state L(eft). Again the underlying mechanism is triggered by the redistribution of non-moving cells upon stimulation which decreases the number of vesicles in C(ortex). If the rates, *κ_gr_* and *κ_cl_*, were equal (i.e. *ν* = 0), then the ratio of L:C (Eq 12) and R:G (Eq 11) would be same before and after stimulation prohibiting upregulation of total transport represented by (*R + L*)/(*G + C*) (Eq (4)). However, with the rate to enter L being smaller than the one to enter R, the decrease in C has a relatively smaller effect on R+L than on G+C and the share of vesicles undergoing directed transport increases.

### Outward directed structural bias of the central transport network promotes overall up-regulation of directed transport upon stimulation

Least squares fitting of the three structural parameters *ρ*_1_, *ρ*_2_ and *ρ*_3_ (Fig 8) by minimising the error functional (7) shows that in order for the model to correctly predict the observed up-regulation of both, outward and total directed transport, the structural parameters *ρ*_1_ and/or *ρ*_2_ are required to have an outward bias (> 0.6). This observation is reminiscent of the requirement that *κ_gr_ > κ_cl_* in the minimal model. It indicates that in the central and sup-peripheral part of the cell the vesicles’ propensity to engage in outward directed transport is higher than to engage in in-ward transport. Note that these parameters are phenomenological, they do not indicate whether this effect is caused by the density of outward-transport fibres in the central and subcortical regions being relatively higher than the density of inward-transport fibres, or – alternatively – by faster rates of attachment to outward directed fibre-motor combinations.

**Fig 8.**
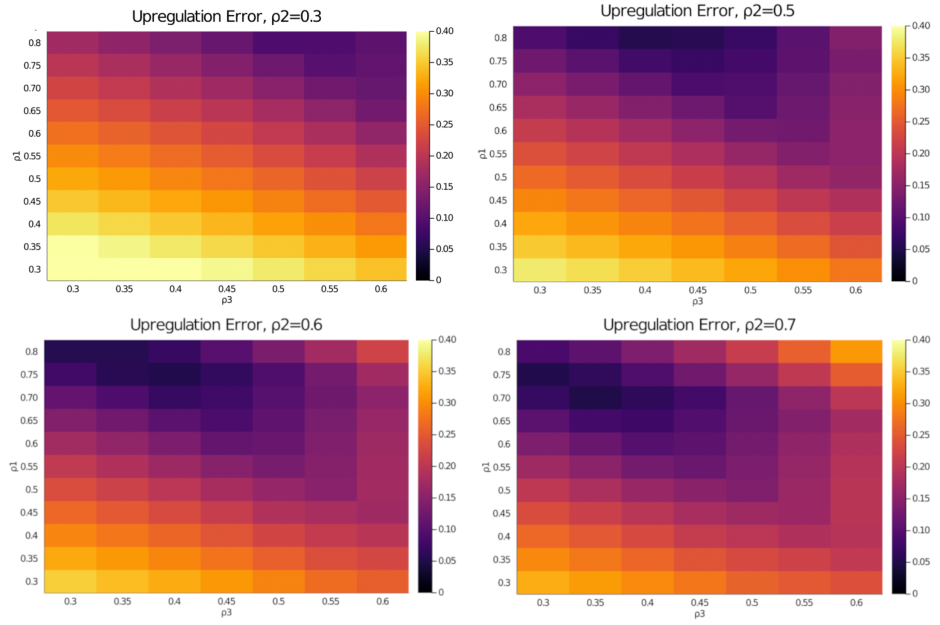
Heatmap plots of the **predictive error** (7) with respect to up-regulation of total and right vs left transport for different values of the structural parameters *ρ*_1_, *ρ*_2_ and *ρ*_3_.

What we conclude is that a cytoskeleton with a bias towards outward transport in the central and subcortical regions will trigger an up-regulation of total and outward vesicle transport in response to their spatial redistribution at stimulation.

### Crowding limits entry into the vesicular transport network towards the plasma membrane

We modelled crowding of cytoskeleton fibres through Hill-functions (8) in the on-rates (9) since a straightforward Markov state model with constant transition rates according to (5) would not be consistent with the reported 3-fold acceleration of transitions of free vesicles into directed motion [19] (reprinted in Fig 2A) upon stimulation.

We fit a single, spatially uniform, repressor coefficient (Table 1) in a way such that - at steady state - the average free-to-directed transition rate is 0.126 min^−1^ in control (0.29 min^−1^ in stimulation) as compared to 0.27 min^−1^ (control) (0.3 min^−1^ in stimulation) for the model with constant transition rates. The model with non-linear transition rates modelling crowding therefore allows to reproduce the 3-fold speed-up of transitions from free into directed motion (Fig 7D) observed experimentally (Fig 2A).

In agreement with the available data (Fig 2B) it predicts that the overall rate of transitions from caged behavior into direction motion does not increase upon stimulation (see Fig 7E). Note that this was only possible through the assumption that crowding doesn’t affect the transitions from caged vesicles in the periphery into directed (see (9)) motion which we believe could reflect the specific structure of the rapidly turning over actomyosin cortex in stimulated cells.

The correction factors with the uniform repressor coefficients listed in Table 1 are close to one in the central compartment, indicating that crowding in control is stronger in the periphery of the cell. (the exact correction factors are visualised in supplementary figure 10). Arguably our choice of the repressor coefficients is simplistic since we lack sufficient data to perform more detailed parameter fitting. It is also for this reason that we don’t perform a detailed sensitivity analysis of the repressor coefficients. Indeed variations in the choice of the repressor coefficients change the magnitude of the correction factors, the only robust feature being that the correction factors are closer to one in the cell centre, whereas crowding is stronger in the periphery. The reason is that most variation of vesicle abundance between control and stimulation is in the periphery of the cell. In the cell centre, however, the model assumes the presence of feedback mechanisms which regulate vesicle synthesis in a way such that vesicle abundance in the cell centre hardly varies between control and stimulation.

## Discussion

In this study we formulate a Markov transition state model which is closely aligned to the data obtained from tracking secretory vesicles in chromaffin cells reported in [19].

We model the asymmetry of directed transport due to the topological constraints of the intracellular space: directed transport originating in the central compartment is biased towards the cell periphery, transport of vesicles originating in the peripheral compartment is towards the cell centre. In the Markov state model Fig 5B this is implemented by the central compartment having only a single transport transition (*R*_1_ → *R*_2_) directed towards the periphery, and by the peripheral compartment having only one transport transition *L*_3_ → *L*_2_ directed towards the cell centre.

Steady state simulations of control and stimulated cells show that secretagogue stimulation shifts the spatial distribution of secretory vesicles from the cell periphery towards the central region of the cell. The spatial redistribution upon stimulation leads to a significant increase of the overall share of vesicles undergoing outward transport. This agrees with observations [19] and with an earlier modelling study [29].

In addition our study suggests that the spatial redistribution also triggers the up-regulation of overall directed transport, i.e. irrespective of direction, observed in [19]. The mechanism relies on the (on-)rates by which vesicles enter the states representing outward transport being faster than the corresponding (on-)rates by which vesicles enter the states associated to inward transport. (Alternatively the (off-)rates by which vesicles leave the states representing outward transport could also be slower than the corresponding rates by which vesicles leave the inward transport states). In this study we illustrate this mechanism using the minimal model visualised in Fig 4.

For the full Markov state model Fig 5 parameter fitting shows that the required asymmetry originates from the central (and - to a lesser extent - the subcortical) compartment having a bias towards outward transport (*ρ*_1_ = 0.75 > 0.5, see Table 1). This indicates that the up-regulation at stimulation is an in-built property of the cell’s cytoskeleton architecture – fibre-motor assemblies in the central region of the cell having a bias towards outward transport – and not necessarily triggered by molecular feedback mechanisms.

Under the additional modelling assumption that the capacity of the cytoskeleton fibres serving as transport tracks is limited in control through crowding, our model also explains the observed accelerated recruitment of free vesicles (vesicles undergoing unconstrained random walks) into directed transport. In which spatial compartment, however, crowding is effective, is difficult to determine based on the published data. The only conclusion we draw is that crowding has a significantly stronger effect in the periphery of the cell as compared to the cell centre, since the steady state distribution of vesicles in control is predominantly characterised by an accumulation of vesicles in the periphery of the cell (see steady state distributions in absolute numbers shown in supplementary Fig 9).

## Conclusion

In this study we show that a mathematical model based on experimentally observed transition rates [19] between motile states of secretory vesicles in chromaffin cells predicts a spatial redistribution of vesicles and up-regulation of outward transport in response to secretagogue stimulation.

We show that this spatial redistribution combined with spatial inhomogeneity of the cellular transport network – a bias towards outward transport in the central region of the cell – explains that upon stimulation a relatively larger share of vesicles undergoes directed transport in agreement with experimental data. This indicates that the up-regulation at stimulation is an in-built property of the cell’s cytoskeleton architecture, and not necessarily triggered by molecular feedback mechanisms.

We also show that crowding of vesicle transport in control, i.e. in non-stimulated cells, is sufficient to explain that the rate by which free vesicles engage in directed transport is three-fold accelerated in stimulated cells [19].

The mathematical model makes a series of testable predictions. While it does not appear feasible to alternate the structure of the cytoskeleton in a way which changes the directional bias of the transport network, manipulation of either plus end or minus end directed molecular motors involved in vesicle transport could have a similar effect. Our model predicts that if the cellular transport network has an inward bias, then the fraction of vesicles undergoing directed transport will be reduced in stimulated cells.

What could be simpler is to limit the rate by which vesicles in stimulated cells progress through the cortical actomyosin network to the site of exocytosis, e.g. through cytoskeleton drugs or genetic manipulation. In this case the Markov state model predicts that the spatial redistribution is reduced and that the up-regulation of both total and outward transport will be limited.

Finally our model also predicts that chromaffin cells with a reduced rate of secretory vesicle biosynthesis will be less affected by crowding in the absence of secretagogue stimulation because of a lower overall vesicle population. Therefore such non-stimulated cells should exhibit the same fast vesicle transition rates from free motion into directed motion as stimulated cells.

One limitation of this study is that it does not consider the details of the pathway by which the actomyosin cortex recruits secretory vesicles and supports their progress towards the active sites. Future research will address this question. Indeed not much is known about the regulation mechanisms that couple cytoplasmic transport of chromaffin vesicles and the regulation of exocytosis through the actin cytoskeleton [11, 18]. We believe that dynamic solutions of mathematical models such as the one introduced in this study have the potential to give insight into those regulation mechanisms, especially when combined with data on vesicle release statistics. Further, they will provide insights for futures studies on other trafficking events including that of autophagosomes [34] and synaptic vesicles in neurons [35].

## Supporting information

### S1 Steady state solution of the minimal 4-state model

The steady state solution of the system (1) satisfies the following equations,

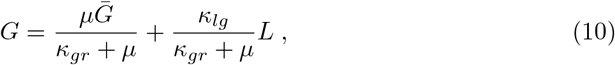

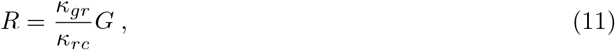

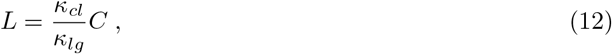

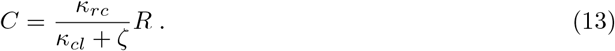

We can now express C and L in terms of G as follows,

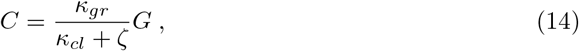

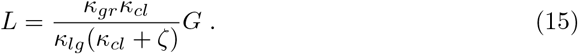

Now we can combine (10), (15) to get the steady state solution for G in terms of 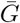,

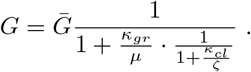

This is the steady state for the population size in state *G*. As mentioned in section Minimal 4-state model it converges to 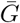 the limit when the rate *μ* is large. This complements the steady state abundances (11),(14) and (15).

### S2 Governing Equations

We provide below the full system of coupled differential equations describing the Markov state model visualised in Figures (5)A and (5)B above using the modified rates from section Modelling of carrying capacities.

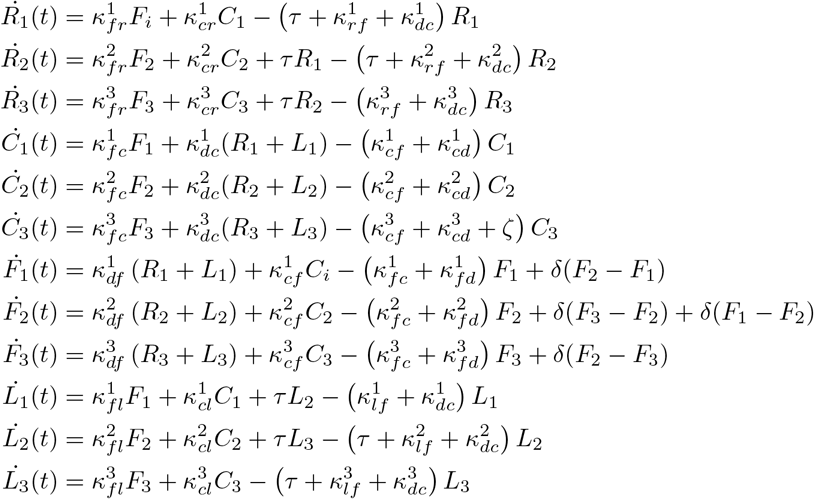

### S3 Steady state distributions, absolute values

**Fig 9.**
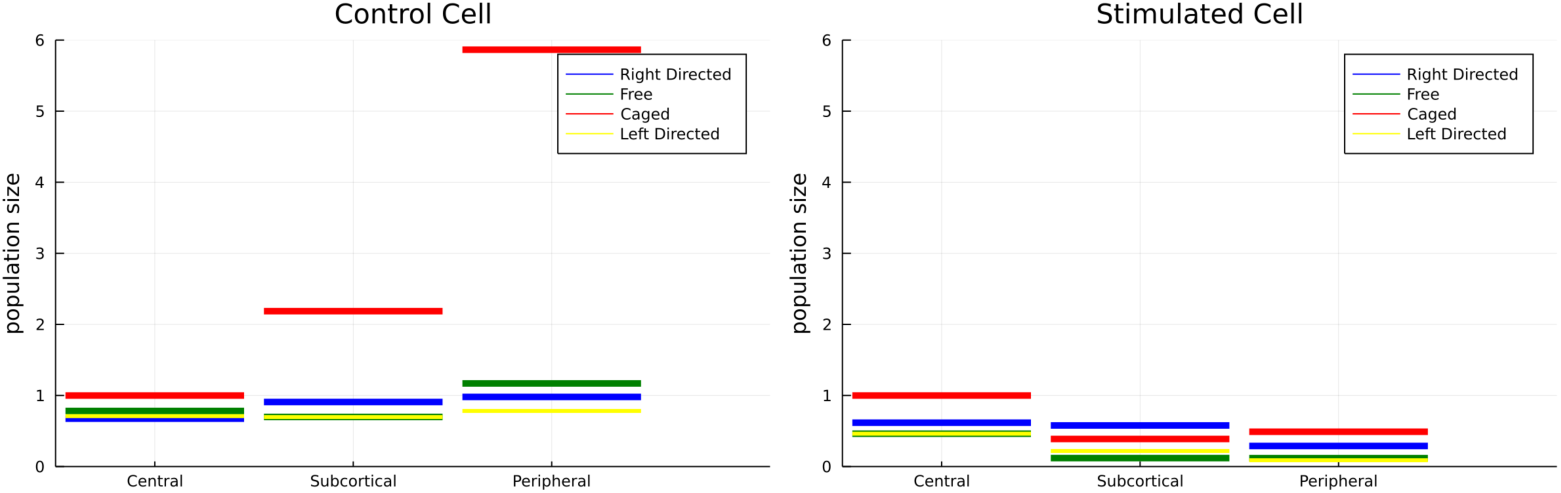
Simulated steady state probability distribution of vesicles in control (A) and upon stimulation (B), visualised with absolute values and *C*1 = 1.

**Fig 10.**
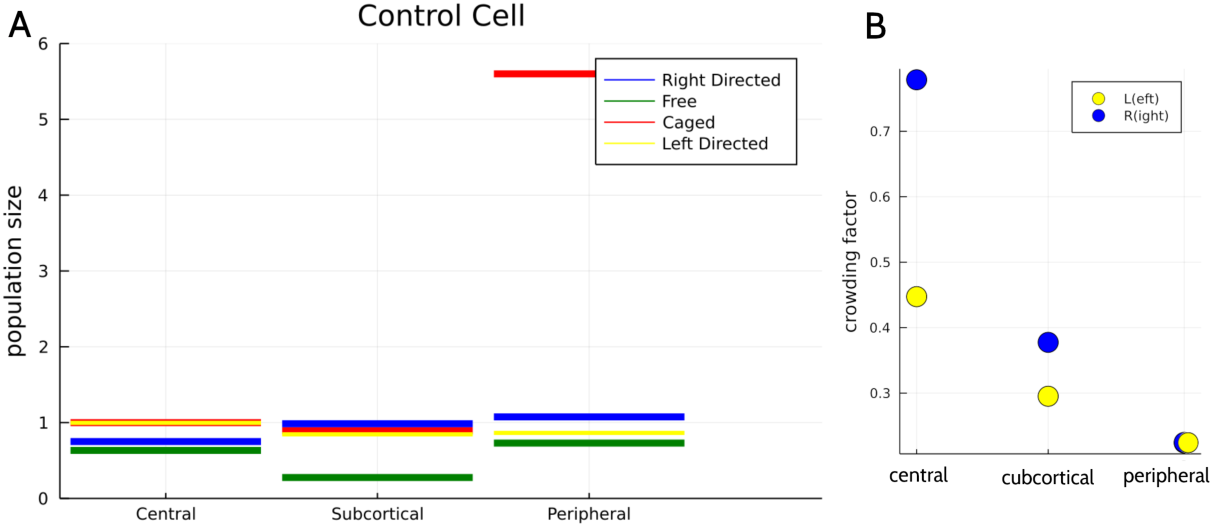
(**A**) Control steady state distribution of vesicles in control with constant transition rates (5) and visualised using absolute values (normalised such that *C*1 = 1). (**B**) Crowding correction factors (8) in control.

## Acknowledgments

DO was supported by Australian Research Council (ARC) Discovery Project DP180102956. FAM was supported by the ARC LIEF grant LE130100078 and by the Australian National Health and Medical Research Council (grants GNT1155794 and GNT1120381 and Senior Research Fellowship 1060075). We used the open-source software Julia [36] with the software packages Catalyst and DifferentialEquations for the numerical computations.

